# Memote: A community driven effort towards a standardized genome-scale metabolic model test suite

**DOI:** 10.1101/350991

**Authors:** Christian Lieven, Moritz E. Beber, Brett G. Olivier, Frank T. Bergmann, Meric Ataman, Parizad Babaei, Jennifer A. Bartell, Lars M. Blank, Siddharth Chauhan, Kevin Correia, Christian Diener, Andreas Dräger, Birgitta E. Ebert, Janaka N. Edirisinghe, Jose P. Faria, Adam Feist, Georgios Fengos, Ronan M. T. Fleming, Beatriz García-Jiménez, Vassily Hatzimanikatis, Wout van Helvoirt, Christopher S. Henry, Henning Hermjakob, Markus J. Herrgård, Hyun Uk Kim, Zachary King, Jasper J. Koehorst, Steffen Klamt, Edda Klipp, Meiyappan Lakshmanan, Nicolas Le Novère, Dong-Yup Lee, Sang Yup Lee, Sunjae Lee, Nathan E. Lewis, Hongwu Ma, Daniel Machado, Radhakrishnan Mahadevan, Paulo Maia, Adil Mardinoglu, Gregory L. Medlock, Jonathan M. Monk, Jens Nielsen, Lars Keld Nielsen, Juan Nogales, Intawat Nookaew, Osbaldo Resendis-Antonio, Bernhard O. Palsson, Jason A. Papin, Kiran R. Patil, Mark Poolman, Nathan D. Price, Anne Richelle, Isabel Rocha, Benjamin J. Sanchez, Peter J. Schaap, Rahuman S. Malik Sheriff, Saeed Shoaie, Nikolaus Sonnenschein, Bas Teusink, Paulo Vilaça, Jon Olav Vik, Judith A. Wodke, Joana C. Xavier, Qianqian Yuan, Maksim Zakhartsev, Cheng Zhang

## Abstract

Several studies have shown that neither the formal representation nor the functional requirements of genome-scale metabolic models (GEMs) are precisely defined. Without a consistent standard, comparability, reproducibility, and interoperability of models across groups and software tools cannot be guaranteed.

Here, we present memote (https://github.com/opencobra/memote) an open-source software containing a community-maintained, standardized set of <u>me</u>tabolic <u>mo</u>del <u>te</u>sts. The tests cover a range of aspects from annotations to conceptual integrity and can be extended to include experimental datasets for automatic model validation. In addition to testing a model once, memote can be configured to do so automatically, i.e., while building a GEM. A comprehensive report displays the model’s performance parameters, which supports informed model development and facilitates error detection.

Memote provides a measure for model quality that is consistent across reconstruction platforms and analysis software and simplifies collaboration within the community by establishing workflows for publicly hosted and version controlled models.

## Introduction

The reconstruction and analysis of metabolic reaction networks provide mechanistic, testable hypotheses for an organism’s metabolism under a wide range of empirical conditions^1^. At the current state of the art, genome-scale metabolic models (GEMs) can include thousands of metabolites and reactions assigned to subcellular locations, gene-protein-reaction rules (GPR), and annotations, which provide meta-information by referencing large biochemical databases. This development has been facilitated by standard protocols for reconstruction^2^ and guidelines for provenance-tracking and interoperability^3–5^. However, the quality control of GEMs remains a formidable challenge that must be solved to enable confident use, reuse, and improvement.

Both Ravikrishnan and Raman^6^ and Ebrahim et al.^7^ lamented the lack of an agreed-upon description format as they found that GEMs can be published as SBML^8^, MATLAB files, spreadsheets, and PDF. While the former noted that incompatible formats limit the scientific exchange and, thus, the ability to reproduce calculations on different setups, the latter elaborated how formatting errors can directly cause inconsistent results when parsed and evaluated with various software packages.

When comparing four previously published models for *Pseudomonas putida* KT2440, Yuan et al. discovered that in identical simulation conditions the predicted growth rate of one model was almost twice as high as that of another^9^. Moreover, one of the examined models could generate ATP without needing to consume any substrate, rendering some model predictions useless.

This behavior occurs when a model’s reaction directions are not checked for thermodynamic feasibility, leading to the formation of flux cycles which provide reduced metabolites to the model without requiring nutrient uptake. Fritzemeier et al.^10^ detected such erroneous energy-generating cycles (EGCs) in the majority of GEMs specifically in the MetaNetX^11,12^ (~66%) and ModelSEED^13^ (~95%) databases, which mostly contain automatically-generated, non-curated metabolic models. Although the authors found that EGCs are rare in manually-curated GEMs from the BiGG Models database (~4%), their effect on the predicted growth rate in FBA may account for an increase of up to 25%. This makes studies involving the growth rates predicted from such models unreliable. It is possible to identify and correct these issues either with functions included in the COBRA Toolbox^14^, or the modified GlobalFit algorithm^15^ presented by Fritzemeier et al.^10^. Yet, as the models of *P. putida* analyzed by Yuan et al. show, this is not done consistently^9^.

Investigating the biomass compositions (BCs) of 71 manually-curated prokaryotic GEMs, Xavier et al. found that organic cofactors (e.g., Coenzyme A, pyridoxal 5-phosphate, and S-adenosyl-methionine) are missing even though their inclusion is vital to a model’s performance in gene-essentiality studies^16^.

Chan et al. highlighted deviations in molecular weight as another problem with the formulation of BCs^17^. Conforming to the defined molecular weight of 1 g/mmol is essential to reliably calculate growth yields, cross-compare models, and obtain valid predictions when simulating microbial consortia. Half of the 64 tested models deviated from the defined 1 g by up to 5%, with the other half differing even more strongly. Any discrepancy, however, should be avoided as the smallest error affects the predicted biomass yield, favoring models containing BCs which sum to lower molecular weight.

In addition to discussing encoding related problems, Ravikrishnan and Raman stressed that missing metabolite and reaction annotations are further fundamental issues when trying to exchange GEMs which have been generated from different platforms, or when attempting to integrate them into existing computational workflows^6^. Mapping annotations between biochemical databases is not trivial but semi-automatic approaches help to reduce the required manual effort^18^. Nonetheless, they reported the absence of metabolite annotations (i.e., metabolite formula, database-dependent (e.g., ChEBI ID), and database-independent i. e. derived from the properties of the object itself (e.g., SMILES, InCHI) references) in almost 60% of the 99 models they examined.

Increasing numbers of manually-curated and automatically-generated GEMs are published each year, growing both in scale and scope; from models on single cells to multi-organism communities^19^ to multi-compartmental plant^20^, human and cancer tissue models^21^. Especially when considering the growing application of models to human health and disease, it becomes essential to address any remaining issues concerning reproducibility and interoperability to pave the way for reliable systems medicine^22^.

Thus, we need to establish a standard framework which ensures that:

- Models are formulated consistently in a software agnostic manner.
- Components of GEMs are uniquely identifiable using standardized database-independent identifiers which can be converted easily using cross-references.
- Default conditions and mathematically specified modeling formulations are precisely defined to allow the reproduction of the original model predictions.
- Models yield biologically feasible phenotypes when analyzed under alternating conditions.
- Data that has been used to curate/parametrize the model are adequately documented to precisely understand the model refinement process.

Here, we argue for a two-pronged approach in creating this framework: 1) We advocate the use of the latest version of the *SBML Level 3 Flux Balance Constraints (FBC) Package^23^* as the agreed-upon description format, which renders GEMs to be independent through a unified formulation. 2) Borrowing tools and best practices from software development^24,25^, we present *memote* as a unified approach for benchmarking metabolic models.

## Results

### SBML: Tool-independent model formulation

Historically, GEMs have been structured and stored in many non-standard ways, for example, tool specific formats or language dialects^6^. This prevented the accurate exchange of models between various software tools and the unambiguous, machine-readable description of all model elements such as chemical reactions, metabolites, gene associations, annotations, objective functions and flux capacity constraints. While a widely used model description standard, such as the Systems Biology Markup Language (SBML) Level 3 Core^8^, can describe some of these components, e.g., reactions, metabolites, or annotations, it cannot present other model components needed to describe a parameterised GEM or FBA model in a structured and semantic way.

Consequently, an adequate model description format is needed that allows for the unambiguous definition and annotation of such a model’s components and underlying mathematics.

With the release of SBML Level 3 it has become possible to load specific modeling packages that extend the core format with additional features. The *SBML Level 3 Flux Balance Constraints (FBC) Package* (SBML3FBC) has been specifically designed to address the problems described above. Such extensions allow users to take advantage of infrastructure built around SBML, while also providing a smaller set of specifications that can be adjusted to cater to the quickly changing needs of a specific research area. The FBC package allows for the unambiguous, tool neutral and validatable SBML description of domain-specific model components such as flux bounds, multiple linear objective functions, gene-protein-reaction associations, metabolite chemical formulas, charge and related annotation^23^. The SBML and constraint-based modeling communities collaboratively develop this package and update it based on user input. As a result, FBC Version 2 is the *de facto* standard for encoding GEMs. Critical to this process is its implementation in a wide range of constraint-based modeling software and adoption by public model repositories^22,26–34^. We believe these factors make SBML3FBC the optimal format for sharing and representing GEMs, thus models encoded in SBML3FBC serve as the input to memote.

### Memote: Community-driven quality control

In software engineering, test-driven development ensures that in response to a defined input a piece of code generates the expected output^35^. Distributed version control represents an efficient way of tracking and merging changes made by a group of people working on the same project^36^. Finally, continuous integration ties these two principles together by automatically triggering tests to be executed after each change that is introduced to the project^37^. Memote (/’mi:moʊt/ (IPA)), short for <u>me</u>tabolic <u>mo</u>del <u>te</u>sts, is an open-source python software that applies these engineering principles to genome-scale metabolic models.

Memote accepts stoichiometric models encoded in SBML3FBC as input, allowing users to benchmark them against a set of consensus tests. By enabling researchers to quickly interrogate the quality of GEMs, problems can be addressed before they affect reproducibility and scientific discourse, or increase the amount of time spent troubleshooting^38^. Memote supports two basic workflows (Figure 1a). First, by running the test suite on a model once, memote generates a comprehensive, human-readable report, which quantifies the model’s performance. By this information, a definitive assessment of model quality can be made, i.e., by editors or reviewers. This workflow is accessible through a web interface (https://memote.dd-decaf.eu), analogous to the SBML validator^27^, or locally through the command line.

Second, for model maintenance and reconstruction, memote coordinates version control and continuous integration, such that each tracked-edit in the reconstruction process can progressively be tested. Users edit the model with their preferred reconstruction tool, and export to SBML afterward. This way, each incremental change can be tested with the suite. Then, a report on the entire history of results serves as a guide towards a functional, high-quality GEM. This workflow is accessible through the command line and may be extended to include custom tests against experimental data. Memote allows researchers to test a model repository offline, but we encourage and support community collaboration in reconstruction via distributed version control development platforms such as GitHub (https://github.com/), GitLab (https://gitlab.com/) or BioModels^39^ (http://wwwdev.ebi.ac.uk/biomodels/).

Either development platform supports a branching strategy (Figure 1b), which model builders could use to curate different parts of the model simultaneously or to invite external experts to improve specific model features. Memote further enables model authors to act as gatekeepers, choosing to accept only high-quality contributions. Identification of functional differences happens in the form of a comparative ‘diff’ report, while for the file-based discrepancies memote capitalizes on the platform’s ability to show the line-by-line changes between different versions of a model. For this purpose, the model is written in a sorted YAML format^40^ after every change.

**Figure 1:**
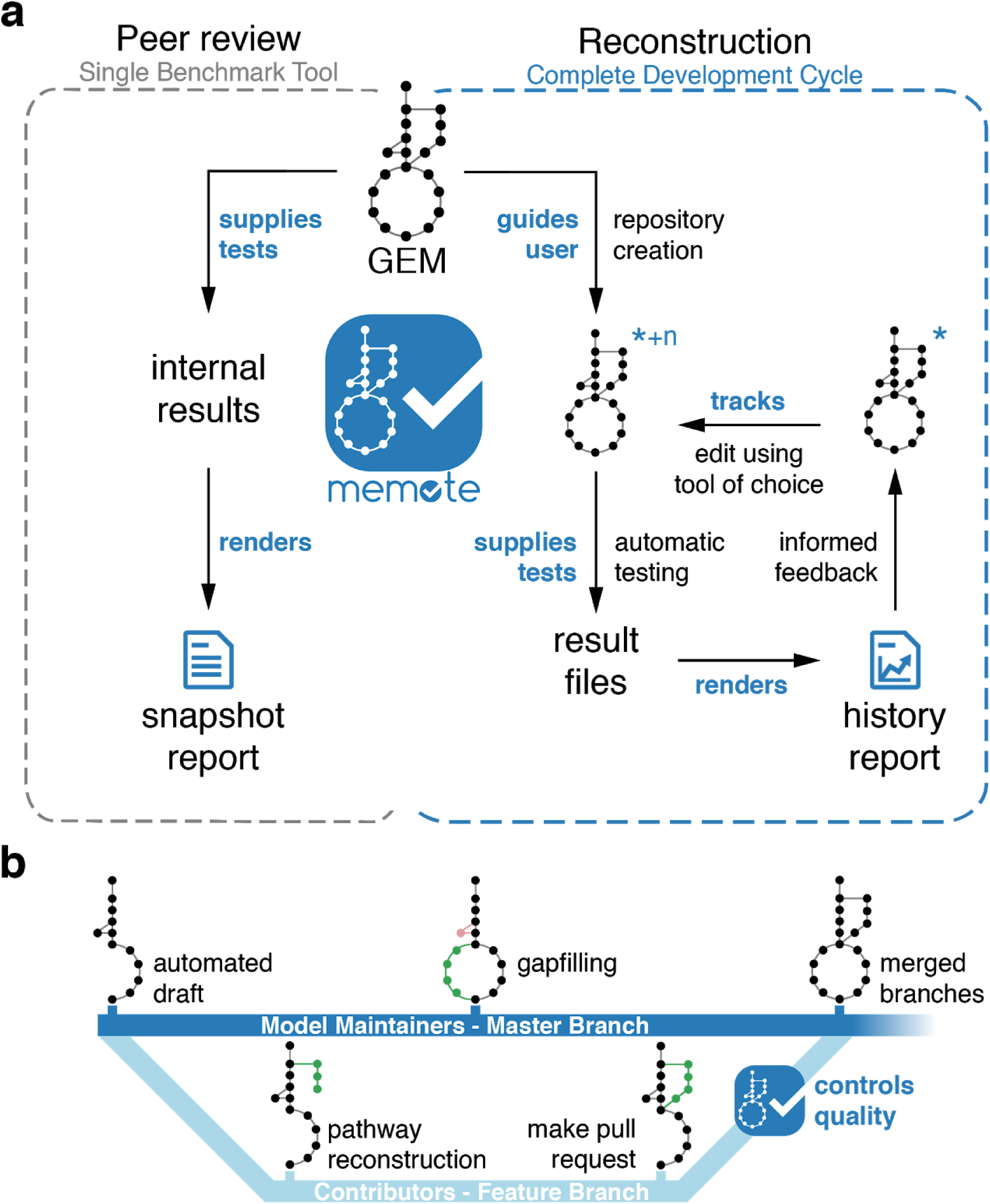
Functionality offered by memote. (a) Graphical representation of the two principal workflows in detail. For peer review, memote serves as a benchmark tool offering a quick snapshot report. For model reconstruction, memote helps the user to create a version-controlled repository for the model (indicated by the **\***), and to activate continuous integration. The model is tested using memote’s library of tests, the results are saved, and an initial report of the model is generated. This constitutes the first iteration of the development cycle. Now, the user may edit the model using a tool of their choice creating a new version (indicated by the **+n**). This will restart the cycle by running the tests automatically, saving the results for each version and including them incrementally in a history report. (b) An example of a potential branching strategy employing memote as a benchmark of external contributions. **Bold blue text** denotes actions performed by memote.

### Description of the test library

The tests within memote are divided into independent core tests and tests that depend on user-supplied experimental data. Core tests are further divided into a scored and an unscored section (Figure 2).

The tests in the scored section are independent of the type of the modeled organism, the complexity of the model itself or the types of identifiers that are used to describe the model components. Calculating a score for these tests allows for the quick comparison of any two given models at a glance. The unscored section provides general statistics and covers specific aspects of a model that are not universally applicable. For instance, dedicated quality control of the biomass equation only applies to metabolic models which are used to investigate cell growth. Tests in either section belong to one of four general areas:

1. Basic tests give an insight into the formal correctness of a model, verifying the existence of the main model components such as metabolites, compartments, reactions, and genes. These tests also check for the presence of formula and charge information of metabolites, and for the presence of gene-protein-reaction rules of reactions. General quality metrics such as the degree of metabolic coverage representing the ratio of reactions and genes^41^ are also covered here.
2. Some tests are dedicated to testing the biomass reaction. This includes testing the model’s ability to produce all biomass precursors in different conditions, the biomass consistency, a non-zero growth rate and direct precursors. The biomass reaction is based on the biomass composition of the modeled organism and expresses its ability to produce the necessary precursors for *in silico* cell growth and maintenance. Hence, an extensive, well-formed biomass reaction is crucial for accurate predictions with a GEM^16^.
3. Stoichiometric inconsistency, erroneously produced energy metabolites^10^ and permanently blocked reactions, are identified by testing the model’s consistency. Errors here may lead to the production of ATP or redox cofactors from nothing^2^ and are detrimental to the performance of the model when using FBA^6^.
4. Annotation tests maintain that a model is equipped according to the community standards with MIRIAM-compliant cross-references^42^, that all primary IDs belong to the same namespace as opposed to being fractured across several namespaces, and that components are described semantically with Systems Biology Ontology terms^43^. A lack of explicit, standardized annotations complicates the use, comparison, and extension of GEMs, and thus strongly hampers collaboration^3,6,44^

**Figure 2:**
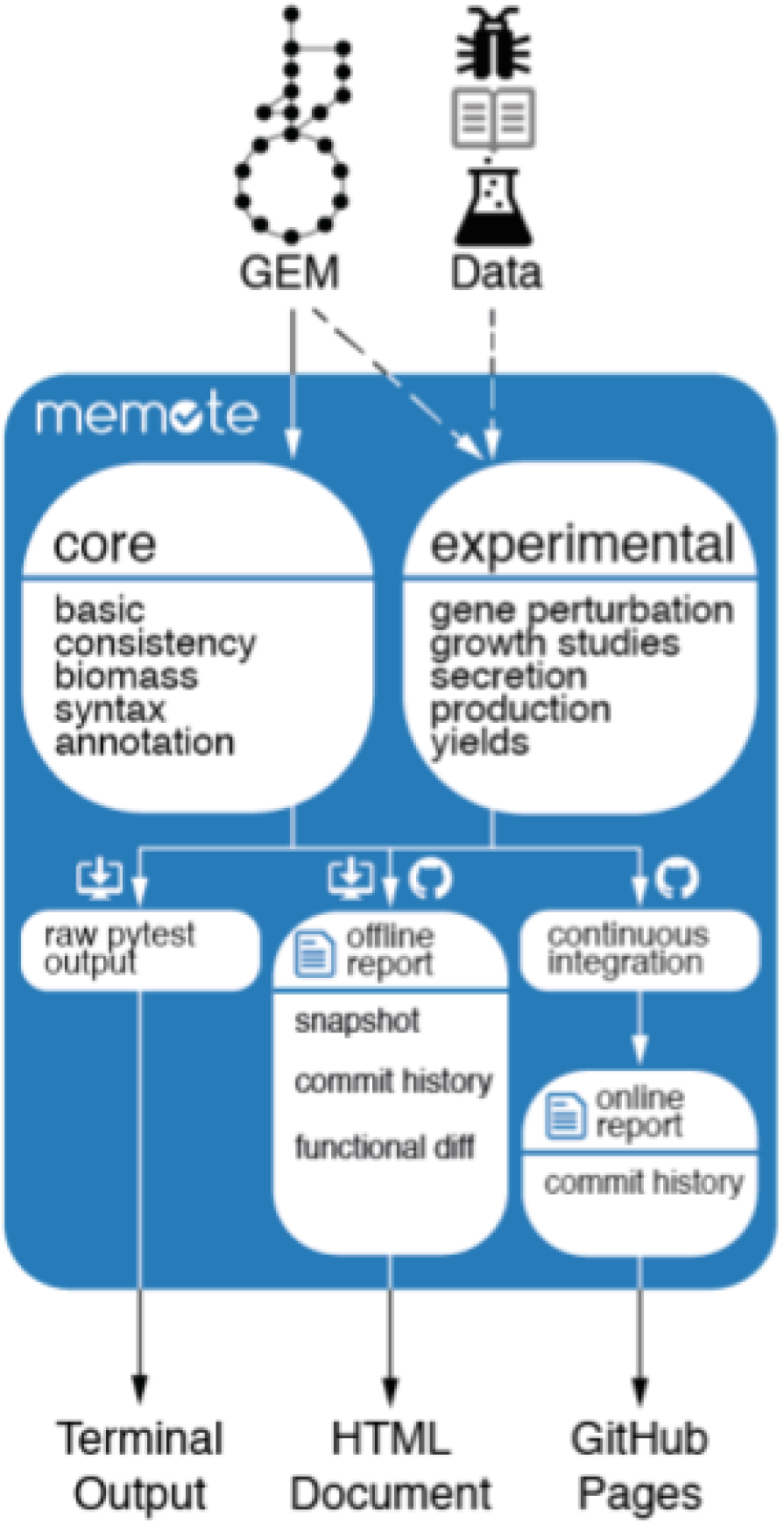
Functional overview of the Metabolic Model Tests (memote) package. A genome-scale metabolic model (GEM) is supplied by the user and tested in all core test categories. Optionally, if the user supplies experimental data, the model will also be subjected to the corresponding experimental tests. After testing, user input through the command line interface determines how the results are displayed. In addition to a high-level output in the terminal, the user can generate a variety of HTML-formatted reports. “Snapshot” will provide a performance benchmark of a single specified model; with a functional diff the user can benchmark two models side-by-side; and the commit-history will show the development of a model’s performance over the course of changes to a version controlled model, The latter is the type of report that is generated automatically when continuous integration is enabled. Then the results are displayed online on the project’s GitHub pages.

A detailed list of all the test in memote is available at https://github.com/opencobra/memote/wiki.

In addition to the core tests, researchers may supply experimental data from gene perturbation studies from a range of input formats (CSV, TSV, XLS or XLSX). Gene perturbation studies, especially gene essentiality studies are useful to refine GEM reconstructions by allowing researchers to identify network gaps and by providing a basis for model validation^45^, as well as providing grounds for a hypothesis about an organism’s physiology^46^.

To constrain the model concerning the experimental conditions underlying the supplied data, researchers may optionally define a configuration file (.yml) in which they can set the medium, FBA objective, and known regulatory effects. Without memote, this would typically be done through the use of custom scripts, which can vary significantly depending on the researcher writing them. Moreover, scripts tend to suffer from software rot if they are not actively maintained after publication^25^. The use of configuration files instead of scripts avoids software rot since the configuration files do not require dependencies other than memote, which is likely to be maintained in the future. In conjunction, setting up a version-controlled model repository not only allows researchers to publish a ‘default’ unspecific GEM of the investigated organism, but also reproducible instructions on how to obtain a model that is specific to the organism in a defined experimental context including, and validated against the data supporting this context. This formulaic approach of deriving a GEM into a condition-specific form supports Heavner and Price’s^3^ call for more transparency and reproducibility in metabolic network reconstruction.

**Figure 3:**
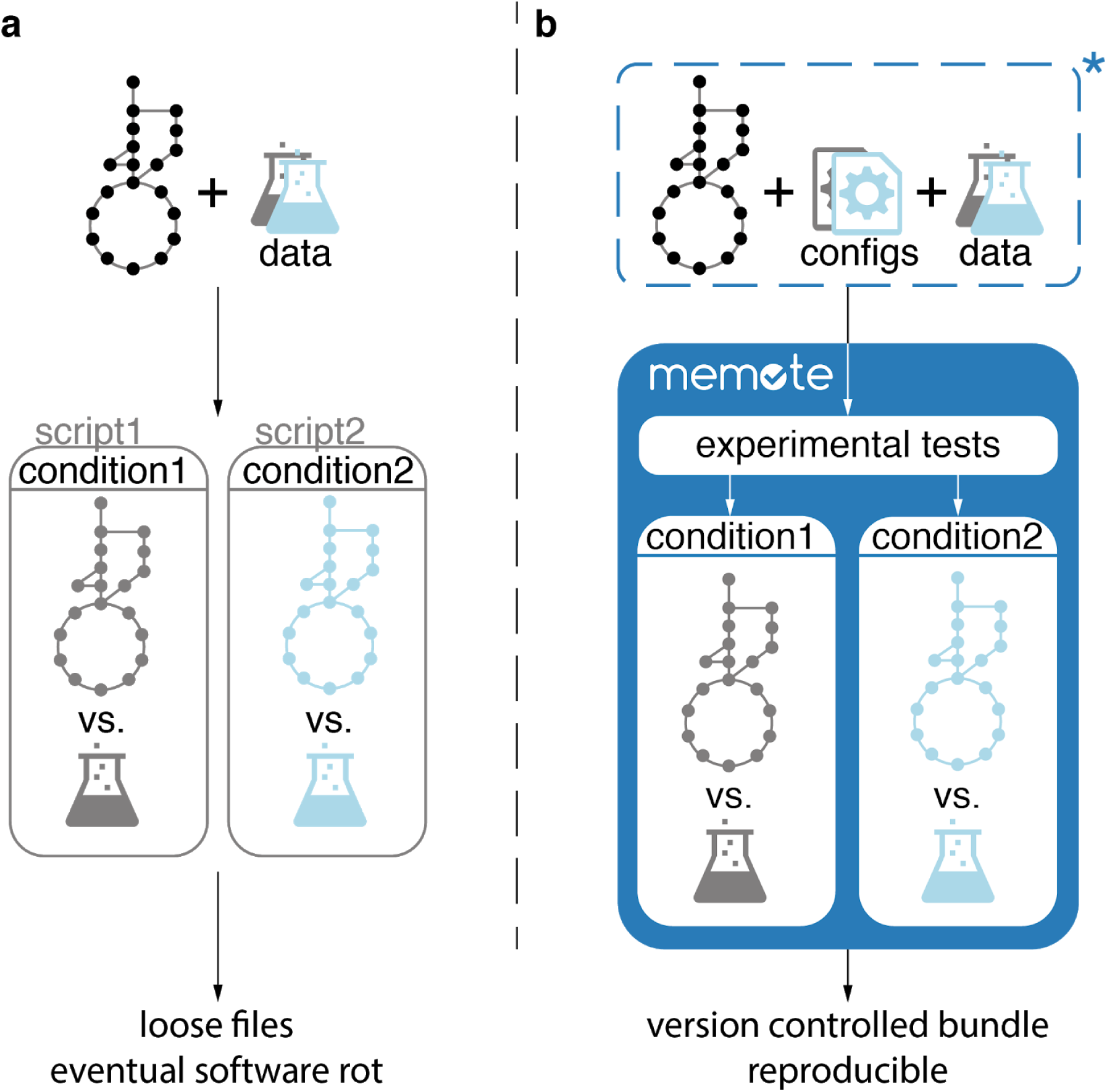
Experimental tests can be tailored to a specific condition through the use of one or several configuration files (configs). (a) To validate GEMs against experimental data measured in specific conditions, researchers usually write their scripts which constrain the model. This is problematic as scripts can vary a lot and they are, unless actively maintained, susceptible to software rot. (b) With memote, user-defined configuration files replace scripts, which allows the experimental validation of GEMs to be unified and formalized. Bundling the model, configuration files, and experimental data within a version-controlled repository (indicated by the **\***) supports cohesive releases.

## Discussion

By providing a performance benchmark based on community guidelines and commonly-referenced SOPs, memote facilitates informed model reconstruction and quality control. The tests within memote cover semantic and conceptual requirements which are fundamental to SBML3FBC and constraint-based modeling, respectively. They are extensible to allow the validation of a model’s performance against experimental data and can be executed as a stand-alone tool or integrated into existing reconstruction pipelines. Capitalizing on robust workflows established in modern software development, memote promotes openness and collaboration by granting the community tangible metrics to support their research and to discuss assumptions or limitations openly.

The concept of having a set of defined metabolic model tests is not dependent on the implementation in memote presented herein. In fact, for some platforms, it may be more desirable to implement these tests separately as this could streamline the user experience. However, an independent, central, community-maintained library of tests and a tool to run them offers 1) an unbiased approach to quality control as the tests are continuously reviewed by the community, 2) a long-lived resource as the project is independent of individual funding sources, 3) flexibility as updates can be propagated rapidly and 4) consistent results as the codebase is unified. To encourage integration as opposed to duplication, memote provides a python API as well as being available as a web-service. We plan to make memote available in the Department of Energy’s Knowledge Base^47^ as an app and integrate it with the BiGG Database^33^, BioModels^31^, and the RAVEN toolbox^48^. The memote test suite plug-in for OptFlux^49^ will approximately be released with version 3.4 scheduled for mid July 2018.

The variety of constraint-based modeling approaches and the fundamental differences between various organisms compound the assessment of GEMs. For instance, authors may publish metabolic networks, which are constrained to reflect one experimental condition or publish unconstrained metabolic databases, which need to be initialized before simulation. Both can be encoded in SBML. With having a scored test section, we attempt to normalize each type of model such that they become comparable. Despite memote’s code itself being unit tested, it is difficult to anticipate all edge cases *a priori*. Also, memote depends on external resources such as MetaNetX^12^ and identifiers.org^50^ that are likely to change over time. Subsequently, individual users may identify potential false-positive and false-negative results. Hence, we recommend to approach the report with scrutiny and encourage users to reach out to the authors to report any errors.

The tests that memote offers only apply to stoichiometric models. However, the underlying principles and individual tests behind memote may apply to models of metabolism and expression (ME-models)^51^, kinetic^52^, or even systems pharmacological models^53^.

The cloud-based distributed version control for GEMs encoded as single SBML files supported by memote is only one possible implementation approach for version control and collaboration on stoichiometric models. For instance, the reconstruction and modeling software Pathway Tools internally stores organism data in the form of a database, which can be queried and altered through the provided guided user interface and access forms^54^ AuReMe, follows a similar approach, by allowing users to interact with a database through automatically generated wikis^55^. While databases offer greater capacity and speed than single, large data files, the programmatic or form-based interaction required for databases may not be most immediately accessible to a broad community.

In the future, with respect to rising big data streams, memote ought to be extended to provide support for tests based on multi-omics data. Moreover, to distribute all files of a model repository together, i.e., the model, supporting data and scripts, these could be automatically bundled into one COMBINE archive file^56^, additionally including SED-ML documents which further describe relevant simulation experiments^57^.

The greater flexibility and awareness of community-driven, open-source development and the trend towards modular approaches exhibited by the solutions that were put forth in the field of systems biology^44^, motivate us to keep the development of memote open. We believe that a robust benchmark can only come to fruition when actively supported by the whole community and thus call for interested experts to involve themselves, be it through testing our tool, discussing its content or improving its implementation. We intend to keep extending memote with additional tests and functionality.

## Acknowledgements

The authors would like to acknowledge Danny Dannaher, Ali Kaafarani and Alba Lopez for their supporting work on the Angular parts of memote; Joao Cardoso, Steinn Gudmundsson, Kristian Jensen and Dimitra Lappa for their feedback on conceptual details; Peter D. Karp and Ines Thiele for critically reviewing the manuscript.

Individual authors acknowledge funding from:

J. O. Vik: the Research Council of Norway grant <u>248792</u> (DigiSal), part of <u>Digital Life Norway</u>.

C. Lieven: Innovation Fund Denmark (project “Environmentally Friendly Protein Production (EFPro2)”).

C. Lieven, N. Sonnenschein, M. Beber, M. Ataman, D. Machado, P. Maia, P. Vilaça, K. R. Patil, and M. Herrgard: This project has received funding from the European Union’s Horizon 2020 research and innovation programme under grant agreement No 686070 (DD-DeCaF). N. E. Lewis: NIGMS R35 GM119850; Novo Nordisk Foundation NNF10CC1016517; Keck Foundation

A. Richelle: Lilly Innovation Fellowship Award

B. García-Jiménez and J. Nogales have received funding from the European Union’s Horizon 2020 research and innovation programme under grant agreement no 686585 for the project LIAR, and the Spanish Ministry of Economy and Competitivity through the RobDcode grant (BIO2014-59528-JIN).

L. M. Blank: The author has received funding from the European Union’s Horizon 2020 research and innovation programme under grant agreement no. 633962 for the project P4SB R. Fleming: U.S. Department of Energy, Offices of Advanced Scientific Computing Research and the Biological and Environmental Research as part of the Scientific Discovery Through Advanced Computing program, grant no. DE-SC0010429.

A. Mardinoglu, C. Zhang, S. Lee and J. Nielsen: The Knut and Alice Wallenberg Foundation. Advanced Computing program, grant #DE-SC0010429.

S. Klamt: This work was in parts supported by the German Federal Ministry of Education and Research (de.NBI partner project “ModSim” (FKZ: 031L104B)).

E. Klipp: This work was supported by the German Federal Ministry of Education and Research (project “SysToxChip”, FKZ 031A303A)

H. U. Kim, S.Y. Lee: Technology Development Program to Solve Climate Changes on Systems Metabolic Engineering for Biorefineries (Grants NRF-2012M1A2A2026556 and NRF-2012M1A2A2026557) from the Ministry of Science and ICT through the National Research Foundation (NRF) of Korea

P. Babaei, Z. King, B. O. Palsson, C. Lieven, M. Beber, N. Sonnenschein, M. Herrgard, A.

Feist: Novo Nordisk Foundation through the Center for Biosustainability at the Technical University of Denmark (NNF10CC1016517)

D.-Y. Lee: Next-Generation BioGreen 21 Program (SSAC, No. PJ01334605), Rural Development Administration, Republic of Korea

G. Fengos: RobustYeast within ERA net project via <u>SystemsX.ch</u>

V. Hatzimanikatis: ETH Domain and Swiss National Science Foundation

M. Poolman: Oxford Brookes University

## Code availability

Memote source code is available at https://github.com/opencobra/memote under the Apache License, Version 2.0.

The supporting documentation ist available at https://memote.readthedocs.io/en/latest/.

The memote web-interface is hosted at https://memote.dd-decaf.eu.

## Author Contributions

CL, MEB and NS conceived the study. MEB, CL, SC and NS wrote the software memote. WvH and JOV alpha-tested memote and provided early ideas for the memote report interface. CL drafted all parts of the manuscript except for the section “SBML: Tool-independent model formulation” which was drafted by BGO and FTB. JOV helped shape the “Introduction”. PM and PV created a memote plug-in for OptFlux. JJK provided a configuration for continuous integration with Gitlab. CL, MEB, BEG, FTB, PB, JAB, LMB, SC, KC, CD, AD, BEE, JNE, AF, RMTF, BGJ, WvH, CSH, HH, MJH, HUK, ZK, JJK, SK, EK, ML, NL, DYL, SYL, SL, NEL, HM, DM, PM, AM, GLM, JMM, RM, JN, LKN, JN, IN, ORA, BØP, JAP, KRP, NDP, AR, IR, PJS, RSMS, SS, NS, BT, PV, JOV, JAW, JCX, QY, MZ, and CZ beta-tested memote and provided feedback and suggestions which shaped the software. All authors read, corrected and approved the manuscript.

## References

1. Palsson, B. Ø. Systems Biology: Constraint-based Reconstruction and Analysis. (Cambridge University Press, 2015).

2. Thiele, I. & Palsson, B. Ø. A protocol for generating a high-quality genome-scale metabolic reconstruction. Nature protocols 5, 93–121 (2010).

3. Heavner, B. D. & Price, N. D. Transparency in metabolic network reconstruction enables scalable biological discovery. Curr. Opin. Biotechnol. 34, 105–109 (2015).

4. Wilkinson, M. D. et al. The FAIR Guiding Principles for scientific data management and stewardship. Sci Data 3, 160018 (2016).

5. van Dam, J. C. J., Koehorst, J. J. J., Vik, J. O., Schaap, P. J. & Suarez-Diez, M. Interoperable genome annotation with GBOL, an extendable infrastructure for functional data mining. *bioRxiv* 184747 (2017). doi:10.1101/184747

6. Ravikrishnan, A. & Raman, K. Critical assessment of genome-scale metabolic networks: the need for a unified standard. Brief. Bioinform. 16, 1057–1068 (2015).

7. Ebrahim, A. et al. Do genome-scale models need exact solvers or clearer standards? Mol. Syst. Biol. 11, 831 (2015).

8. Hucka, M. et al. The Systems Biology Markup Language (SBML): Language Specification for Level 3 Version 1 Core. 167 (2010).

9. Yuan, Q. et al. Pathway-Consensus Approach to Metabolic Network Reconstruction for Pseudomonas putida KT2440 by Systematic Comparison of Published Models. PLoS One 12, e0169437 (2017).

10. Fritzemeier, C. J., Hartleb, D., Szappanos, B., Papp, B. & Lercher, M. J. Erroneous energy-generating cycles in published genome scale metabolic networks: Identification and removal. PLoS Comput. Biol. 13, e1005494 (2017).

11. Ganter, M., Bernard, T., Moretti, S., Stelling, J. & Pagni, M. MetaNetX.org: a website and repository for accessing, analysing and manipulating metabolic networks. Bioinformatics 29, 815–816 (2013).

12. Moretti, S. et al. MetaNetX/MNXref-reconciliation of metabolites and biochemical reactions to bring together genome-scale metabolic networks. Nucleic Acids Res. 44, D523–D526 (2016).

13. Henry, C. S. et al. High-throughput generation, optimization and analysis of genome-scale metabolic models. Nat. Biotechnol. 28, 977–982 (2010).

14. Heirendt, L. et al. Creation and analysis of biochemical constraint-based models: the COBRA Toolbox v3.0. *arXiv [q-bio.QM]* (2017).

15. Hartleb, D., Jarre, F. & Lercher, M. J. Improved Metabolic Models for E. coli and Mycoplasma genitalium from GlobalFit, an Algorithm That Simultaneously Matches Growth and Non-Growth Data Sets. PLoS Comput. Biol. 12, e1005036 (2016).

16. Xavier, J. C., Patil, K. R. & Rocha, I. Integration of Biomass Formulations of Genome-Scale Metabolic Models with Experimental Data Reveals Universally Essential Cofactors in Prokaryotes. Metab. Eng. 39, 200–208 (2017).

17. Chan, S. H. J., Cai, J., Wang, L., Simons-Senftle, M. N. & Maranas, C. D. Standardizing biomass reactions and ensuring complete mass balance in genome-scale metabolic models. Bioinformatics (2017). doi:10.1093/bioinformatics/btx453

18. Haraldsdóttir, H. S., Thiele, I. & Fleming, R. M. Comparative evaluation of open source software for mapping between metabolite identifiers in metabolic network reconstructions: application to Recon 2. J. Cheminform. 6, 2 (2014).

19. Kim, W. J., Kim, H. U. & Lee, S. Y. Current state and applications of microbial genome-scale metabolic models. Current Opinion in Systems Biology 2, 10–18 (2017).

20. Zakhartsev, M. et al. Metabolic model of central carbon and energy metabolisms of growing Arabidopsis thaliana in relation to sucrose translocation. BMC Plant Biol. 16, 262 (2016).

21. Jerby, L. & Ruppin, E. Predicting drug targets and biomarkers of cancer via genome-scale metabolic modeling. Clin. Cancer Res. 18, 5572–5584 (2012).

22. Olivier, B. G., Swat, M. J. & Moné, M. J. Modeling and Simulation Tools: From Systems Biology to Systems Medicine. Methods Mol. Biol. 1386, 441–463 (2016).

23. Olivier, B. G. & Bergmann, F. T. SBML Level 3 Package: Flux Balance Constraints version 2. J. Integr. Bioinform. 15, (2018).

24. Cooper, J., Vik, J. O. & Waltemath, D. A call for virtual experiments: accelerating the scientific process. Prog. Biophys. Mol. Biol. 117, 99–106 (2015).

25. Beaulieu-Jones, B. K. & Greene, C. S. Reproducibility of computational workflows is automated using continuous analysis. Nat. Biotechnol. 35, 342–346 (2017).

26. Bergmann, F. T. & Sauro, H. M. SBW - A Modular Framework for Systems Biology. in Proceedings of the 2006 Winter Simulation Conference 1637–1645 (2006). doi: 10.1109/WSC.2006.322938

27. Bornstein, B. J., Keating, S. M., Jouraku, A. & Hucka, M. LibSBML: an API library for SBML. Bioinformatics 24, 880–881 (2008).

28. Olivier, B. G. PySCeS CBMPy: Constraint Based Modelling in Python. (2011).

29. Frank T. Bergmann, S. M. K. SBML Team facilities & software. (2011). doi:10.1038/npre.2011.6401. 1

30. Ebrahim, A., Lerman, J. A., Palsson, B. O. & Hyduke, D. R. COBRApy: COnstraints-Based Reconstruction and Analysis for Python. BMC Syst. Biol. 7, 74 (2013).

31. Chelliah, V. et al. BioModels: ten-year anniversary. Nucleic Acids Res. 43, D542–8 (2015).

32. Rodriguez, N. et al. JSBML 1.0: providing a smorgasbord of options to encode systems biology models. Bioinformatics 31, 3383–3386 (2015).

33. King, Z. A. et al. BiGG Models: A platform for integrating, standardizing and sharing genome-scale models. Nucleic Acids Res. 44, D515–22 (2016).

34. Cardoso, J., Jensen, K., Lieven, C. & Hansen, A. Cameo: A Python Library for Computer Aided Metabolic Engineering and Optimization of Cell Factories. *bioRxiv* (2017).

35. Nagappan, N., Michael Maximilien, E., Bhat, T. & Williams, L. Realizing quality improvement through test driven development: results and experiences of four industrial teams. Empir. Softw. Eng. 13, 289–302 (2008).

36. Brindescu, C., Codoban, M., Shmarkatiuk, S. & Dig, D. How Do Centralized and Distributed Version Control Systems Impact Software Changes? in Proceedings of the 36th International Conference on Software Engineering 322–333 (ACM, 2014). doi: 10.1145/2568225.2568322

37. Meyer, M. Continuous Integration and Its Tools. IEEE Softw. 31, 14–16 (2014).

38. Baker, M. How quality control could save your science. Nature 529, 456–458 (2016).

39. Glont, M. et al. BioModels: expanding horizons to include more modelling approaches and formats. Nucleic Acids Res. 46, D1248–D1253 (2018).

40. Steffensen, J. L., Dufault-Thompson, K. & Zhang, Y. PSAMM: A Portable System for the Analysis of Metabolic Models. PLoS Comput. Biol. 12, e1004732 (2016).

41. Monk, J., Nogales, J. & Palsson, B. O. Optimizing genome-scale network reconstructions. Nat. Biotechnol. 32, 447–452 (2014).

42. Le Novére, N. et al. Minimum information requested in the annotation of biochemical models (MIRIAM). Nat. Biotechnol. 23, 1509–1515 (2005).

43. Courtot, M. et al. Controlled vocabularies and semantics in systems biology. Mol. Syst. Biol. 7, 543 (2011).

44. Dräger, A. & Palsson, B. Ø. Improving collaboration by standardization efforts in systems biology. Front Bioeng Biotechnol 2, 61 (2014).

45. Feist, A. M., Herrgard, M. J., Thiele, I., Reed, J. L. & Palsson, B. O. Reconstruction of Biochemical Networks in Microbial Organisms. Nat. Rev. Microbiol. 7, 129–143 (2009).

46. Oberhardt, M. A., Palsson, B. Ø. & Papin, J. A. Applications of genome-scale metabolic reconstructions. Mol. Syst. Biol. 5, 320 (2009).

47. Arkin, A. P. et al. The DOE Systems Biology Knowledgebase (KBase). *bioRxiv* 096354 (2016). doi:10.1101/096354

48. Agren, R. et al. The RAVEN toolbox and its use for generating a genome-scale metabolic model for Penicillium chrysogenum. PLoS Comput. Biol. 9, e1002980 (2013).

49. Rocha, I. et al. OptFlux: an open-source software platform for in silico metabolic engineering. BMC Syst. Biol. 4, 45 (2010).

50. Juty, N., Le Novère, N. & Laibe, C. Identifiers.org and MIRIAM Registry: community resources to provide persistent identification. Nucleic Acids Res. 40, D580–6 (2012).

51. O’Brien, E. J., Lerman, J. A., Chang, R. L., Hyduke, D. R. & Palsson, B. Ø. Genome-scale models of metabolism and gene expression extend and refine growth phenotype prediction. Mol. Syst. Biol. 9, 693 (2013).

52. Vasilakou, E. et al. Current state and challenges for dynamic metabolic modeling. Curr. Opin. Microbiol. 33, 97–104 (2016).

53. Thiel, C. et al. A Comparative Analysis of Drug-Induced Hepatotoxicity in Clinically Relevant Situations. PLoS Comput. Biol. 13, e1005280 (2017).

54. Karp, P. D. et al. Pathway Tools version 13.0: integrated software for pathway/genome informatics and systems biology. Brief. Bioinform. 11, 40–79 (2010).

55. Aite, M. et al. Traceability, reproducibility and wiki-exploration for ‘à-la-carte’ reconstructions of genome-scale metabolic models. PLoS Comput. Biol. 14, e1006146 (2018).

56. Bergmann, F. T. et al. COMBINE archive and OMEX format: one file to share all information to reproduce a modeling project. BMC Bioinformatics 15, 369 (2014).

57. Waltemath, D. et al. Reproducible computational biology experiments with SED-ML-the Simulation Experiment Description Markup Language. BMC Syst. Biol. 5, 198 (2011).

